# Humoral immunity induced by LP.8.1 monovalent vaccines against a broad range of SARS-CoV-2 variants including XEC, NB.1.8.1, XFG, and BA.3.2

**DOI:** 10.1101/2025.11.18.689152

**Authors:** Yu Kaku, Mizuka Fujiwara, Keiya Uriu, Maximilian Stanley Yo, Shusuke Kawakubo, Jumpei Ito, Naoya Itoh, Yoshifumi Uwamino, Fumitake Saito, Hironori Satoh, The Genotype to Phenotype Japan (G2P-Japan) Consortium, Kei Sato

## Abstract

In the spring of 2025, multiple SARS-CoV-2 Omicron JN.1 subvariants were circulating, with LP.8.1 among the major variants. Pharmaceutical companies such as Pfizer/BioNTech, Moderna, and Novavax/Takeda adopted monovalent LP.8.1 for their 2025–2026 season vaccines, following recommendations issued by the WHO in May 2025. As of November 2025, SARS-CoV-2 variants including LP.8.1, XEC, NB.1.8.1, and XFG—all designated as variants under monitoring—were circulating. In terms of the spike gene, these recent variants as well as LP.8.1 are derived from JN.1. Moreover, BA.3.2, a BA.3 descendant with multiple mutations in the spike gene, has recently emerged and exhibits robust immune evasion. In Japan, the rollout of the LP.8.1-based vaccination has progressed since the end of September 2025. We previously reported the humoral immunity induced by the XBB.1.5-based monovalent vaccine in 2023 and the JN.1-based monovalent vaccine in 2024 in the Japanese population. Here, we investigated the efficiency of humoral immunity induced by two LP.8.1-based vaccines, the mRNA vaccine from Pfizer/BioNTech and the recombinant protein-based vaccine from Novavax/Takeda, in Japan. We performed neutralization assays using sera obtained from individuals who received the LP.8.1 mRNA vaccine from Pfizer/BioNTech (N=29) or the LP.8.1 recombinant protein vaccine from Novavax/Takeda (N=20) with pseudoviruses harboring spike proteins of B.1.1, BA.5, XBB.1.5, JN.1, LP.8.1, XEC, NB.1.8.1, XFG and BA.3.2. In both mRNA and protein-based vaccinee groups, the change in 50% neutralizing titer (NT50) against variants that were predominant before JN.1 (i.e., B.1.1, BA.5 and XBB.1.5) were smaller than those against JN.1 and its subvariants, including LP.8.1, XEC, NB.1.8.1 and XFG. Consistent with recent studies, neutralizing antibodies against BA.3.2 were induced by both vaccines. However, the induction fold change of BA.3.2 was smaller than those of JN.1 and its subvariants. Next, we tested humoral immune response of participants who received both JN.1-based vaccine in 2024 and LP.8.1-based vaccine in 2025 (N=15). Approximately a year after the JN.1-based vaccination, neutralization titer has waned against all variants tested. However, when we compare the NT50s of pre-vaccination sera between 2025 and 2024, those in 2025 against all variants except for B.1.1 were significantly higher than those last year. This suggests that the cross-neutralizing antibodies induced by JN.1-based vaccination were still maintained for a year. Furthermore, the neutralization abilities against the JN.1 sublineages tested and BA.3.2 were significantly reboosted after the LP.8.1-based vaccination. Our study shows immune boosting by the LP.8.1-based vaccine is effective in achieving cross-neutralization against a broad range of JN.1 sublineages in a JN.1-naïve population and in recalling waning humoral immunity against these subvariants.

## Text

In the spring of 2025, multiple SARS-CoV-2 Omicron JN.1 subvariants were circulating, with LP.8.1 among the major variants. Pharmaceutical companies such as Pfizer/BioNTech, Moderna, and Novavax/Takeda adopted monovalent LP.8.1 for their 2025–2026 season vaccines, following recommendations issued by the WHO in May 2025 (https://www.who.int/news/item/15-05-2025-statement-on-the-antigen-composition-of-covid-19-vaccines). As of November 2025, SARS-CoV-2 variants including LP.8.1, XEC, NB.1.8.1, and XFG—all designated as variants under monitoring—were circulating.^1^ In terms of the spike gene, these recent variants as well as LP.8.1 are derived from JN.1. Moreover, BA.3.2, a BA.3 descendant with multiple mutations in the spike gene, has recently emerged and exhibits robust immune evasion^2,3^. In Japan, the rollout of the LP.8.1-based vaccination has progressed since the end of September 2025. We previously reported the humoral immunity induced by the XBB.1.5-based monovalent vaccine in 2023^4^ and the JN.1-based monovalent vaccine in 2024^5,6^ in the Japanese population. Here, we investigated the efficiency of humoral immunity induced by two LP.8.1-based vaccines, the mRNA vaccine from Pfizer/BioNTech and the recombinant protein-based vaccine from Novavax/Takeda, in Japan.

We performed neutralization assays using sera obtained from individuals who received the LP.8.1 mRNA vaccine from Pfizer/BioNTech (N=29; **Figure A**) or the LP.8.1 recombinant protein vaccine from Novavax/Takeda (N=20; **Figure B**) with pseudoviruses harboring spike proteins of B.1.1, BA.5, XBB.1.5, JN.1, LP.8.1, XEC, NB.1.8.1, XFG and BA.3.2. In both mRNA and protein-based vaccinee groups, the change in 50% neutralizing titer (NT50) against variants that were predominant before JN.1 (i.e., B.1.1, BA.5 and XBB.1.5) were smaller than those against JN.1 and its subvariants, including LP.8.1, XEC, NB.1.8.1 and XFG. Consistent with recent studies^7,8^, neutralizing antibodies against BA.3.2 were induced by both vaccines. However, the induction fold change of BA.3.2 was smaller than those of JN.1 and its subvariants.

**Figure.**
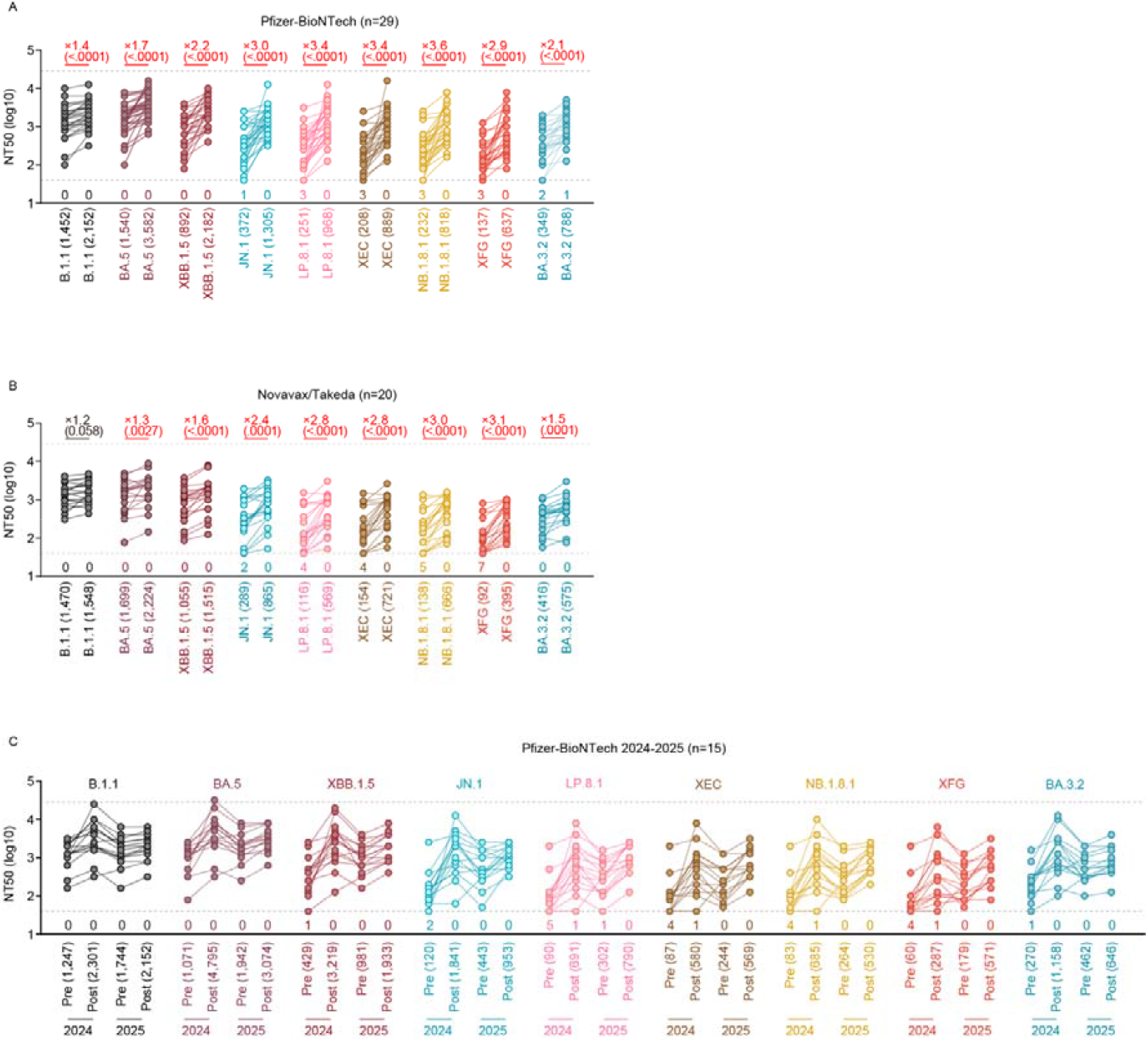
The cross-neutralization induced by LP.8.1 monovalent vaccine. Neutralization assay. Assays were performed with pseudoviruses harboring the spike proteins of B.1.1, BA.5, XBB.1.5, JN.1, LP.8.1, XEC, NB.1.8.1, XFG and BA.3.2. The following two sera were used: LP.8.1 monovalent vaccine sera from fully vaccinated individuals who had received Pfizer-BioNTech LP.8.1 vaccine (“Pfizer-BioNTech” cohort) (three 3-dose vaccinated, nine 4-dose vaccinated, six 5-dose vaccinated, five 6-dose vaccinated, three 7-dose vaccinated and three 8-dose vaccinated; twenty nine donors, average age: 33.8, range: 19–56, 37.9% male) (**A**), and those from fully vaccinated individuals who received Novavax/Takeda LP.8.1 vaccine (“Novavax/Takeda “ cohort) (two 5-dose vaccinated, eleven 6-dose vaccinated, five 7-dose vaccinated, one 8-dose vaccinated and one 9-dose vaccinated; twenty donors, average age: 44.5, range: 23-58, 45.0% male) (**B**). Assays were also performed using sera from fully vaccinated individuals who had received and LP.8.1 vaccine (included in “Pfizer-BioNTech” cohort) (three 3-dose vaccinated, two 4-dose vaccinated, four 5-dose vaccinated, three 6-dose vaccinated and three 8-dose vaccinated; fifteen nine donors, average age: 35.3, range: 25–56, 53.3% male) and sera from the same individuals when they received Pfizer-BioNTech JN.1 vaccine (**C**). Assays for each serum sample were performed in quadruplicate to determine the 50% neutralization titer (NT50). Each dot represents one NT50 value for each donor, and the NT50 values for the same donor before and after vaccination are connected by a line. Numbers in parentheses below the graphs indicate the median of the NT50 value. The horizontal dashed lines indicate the detection limit of the lowest (40-fold dilution) and the highest (29,160-fold dilution), respectively. The number of sera with the NT50 values below the lower detection limit is shown in the figure (under the bottom horizontal dashed line). Neutralization titers below the lower and above the higher detection limit were calculated as titers of 40 and 29,160 (**A-C**). The fold changes of NT50 between before and after vaccination indicated in red with “X” are calculated as the median of ratio of NT50 obtained from each individual. The p-values in the parentheses were determined by two-sided Wilcoxon signed-rank tests. The NT50s against each virus are displayed with sera collected before vaccination on the left and sera collected 3-4 weeks after vaccination on the right of each block on the X-axis (**A-B**). These also indicated as ‘Pre’ and ‘Post’ from left to right in 2024 and 2025 (**C**).

Next, we tested humoral immune response of participants who received both JN.1-based vaccine in 2024 and LP.8.1-based vaccine in 2025 (N=15, **Figure C**). Approximately a year after the JN.1-based vaccination, neutralization titer has waned (0.3-fold to 0.6-fold, *P*<0.02) against all variants tested. However, when we compare the NT50s of pre-vaccination sera between 2025 and 2024, those in 2025 against all variants except for B.1.1 were significantly higher than those last year (**Figure C**). This suggests that the cross-neutralizing antibodies induced by JN.1-based vaccination were still maintained for a year. Furthermore, the neutralization abilities against the JN.1 sublineages tested and BA.3.2 were significantly reboosted after the LP.8.1-based vaccination (**Figure C**). Our study shows immune boosting by the LP.8.1-based vaccine is effective in achieving cross-neutralization against a broad range of JN.1 sublineages in a JN.1-naïve population and in recalling waning humoral immunity against these subvariants.

## References

1. World Health Organization, WHO COVID-19 dashboard, https://data.who.int/dashboards/covid19/variants, Date: 2025, Date accessed: November 16, 2025.

2. Zhang L, Chen N, Eichmann A, et al. Epidemiological and virological update on the emerging SARS-CoV-2 variant BA.3.2. Lancet Infect Dis 2025.

3. Guo C, Yu Y, Liu J, et al. Antigenic and virological characteristics of SARS-CoV-2 variants BA.3.2, XFG, and NB.1.8.1. Lancet Infect Dis 2025; 25:7e374–e377.

4. Kosugi Y, Kaku Y, Hinay AAJ, et al. Antiviral humoral immunity against SARS-CoV-2 omicron subvariants induced by XBB.1.5 monovalent vaccine in infectionnaive and XBB-infected individuals. Lancet Infect Dis 2024; 24(3): e147–8.

5. Uriu K, Kaku Y, Uwamino Y, et al. Antiviral humoral immunity induced by JN.1 monovalent mRNA vaccines against SARS-CoV-2 omicron subvariants including JN.1, KP.3.1.1, and XEC. Lancet Infect Dis 2025; 25(2):e61.

6. Uriu K, Kaku Y, Kihara M, et al. Antiviral humoral immunity against SARS-CoV-2 omicron JN.1 subvariants induced by JN.1 recombinant protein vaccine. Vaccine 2025; 30:54:127129.

7. Nasir A, Lee DW, Wang Z, et al. mRNA-1273.251 and mRNA-1283.251 vaccines expressing SARS-CoV-2 variant LP.8.1 antigens broadly neutralize contemporary JN.1-lineage viruses. bioRxiv. 2025. doi: 10.1101/2025.09.16.676634

8. Hoffmann HM, Stankov MV, Nehlmeier I, et al. Impact of LP.8.1-Adapted mRNA Vaccination on SARS-CoV-2 Variant Neutralisation. medRxiv 2025. doi: 10.1101/2025.10.21.25338461.

