## Appendix for "Humoral immunity induced by LP.8.1 monovalent vaccines against a broad range of SARS-CoV-2 variants including XEC, NB.1.8.1, XFG, and BA.3.2"

**Supplementary Appendix**

**Table of Contents**

**Contents** Page

**Materials and Methods** 2-3
Ethics statement
Human serum collection
Cell culture
Pseudovirus preparation
Neutralization assay
**Table S1.** Human vaccination sera used in this study 4
**Table S2.** Primers in this study 5
**Table S3.** Statistics for neutralization titers in 2024 and 2025 6
**Consortia** 7-8
**Acknowledgments** 9
**Supplemental References** 10

### Materials and Methods

#### Ethics statement

All protocols involving specimens from human subjects recruited at Keio University, Sato naika clinic, and Namikibashi Clinic were reviewed and approved by the Institutional Review Boards of The Institute of Medical Science, The University of Tokyo (approval IDs: 2021-1-0416 and 2022-29-0915), and Keio University (approval ID: 20200059), respectively. All human subjects provided written informed consent. All protocols for the use of human specimens were reviewed and approved by the Institutional Review Boards of The Institute of Medical Science, The University of Tokyo (approval IDs: 2021-1-0416, 2021-18-0617 and 2022-29-0915).

#### Human serum collection

LP.8.1 monovalent vaccine sera from fully vaccinated individuals who had received Pfizer-BioNTech LP.8.1 vaccine ("Pfizer-BioNTech" cohort) (three 3-dose vaccinated, nine 4-dose vaccinated, six 5-dose vaccinated, five 6-dose vaccinated, three 7-dose vaccinated and three 8-dose vaccinated; twenty nine donors, average age: 33.8, range: 19–56, 37.9% male) and those from fully vaccinated individuals who received Novavax/Takeda LP.8.1 vaccine ("Novavax/Takeda" cohort) (two 5-dose vaccinated, eleven 6-dose vaccinated, five 7-dose vaccinated, one 8-dose vaccinated and one 9-dose vaccinated; twenty donors, average age: 44.5, range: 23-58, 45.0% male). For this study, we collected samples before vaccination and three to four weeks (21–29 days) after vaccination.
Sera were inactivated at 56°C for 30 minutes and stored at –80°C until use. The details of the convalescent sera are summarized in **Table S1**.

#### Cell culture

The Lenti-X 293T cells (Takara, Cat# 632180) and HOS-ACE2/TMPRSS2 cells (kindly provided by Dr. Kenzo Tokunaga), a derivative of HOS cells (a human osteosarcoma cell line; ATCC CRL-1543) stably expressing human ACE2 and TMPRSS2[1,2] were maintained in Dulbecco's modified Eagle's medium (DMEM) (high glucose) (Wako, Cat# 044- 29765) containing 10% fetal bovine serum (Sigma-Aldrich Cat# 172012-500ML), 100 units penicillin and 100 ug/ml streptomycin (Sigma-Aldrich, Cat# P4333-100ML).

#### Pseudovirus preparation

Plasmids expressing the SARS-CoV-2 spike (S) proteins of B.1.1, BA.5, XBB.1.5, JN.1, LP.8.1, XEC and NB.1.8.1 were prepared in our previous studies.[3-10] A plasmid expressing the XFG spike protein was generated by site-directed overlap extension PCR using pC-SARS2-S NB.1.8.1 as the template and primers listed in **Table S2**. An oligonucleotide of the BA.3.2 spike protein was synthesized by FACSMAC DNA Synthesis service (https://fasmac.co.jp/en/dnasynthesis) based on the sequence deposited in GISAID (Accession ID: EPI_ISL_19893762). This sequence, belonging to the BA.3.2.2 sublineage, was selected as reference because it contains no undetermined amino acid 'X' residues in the spike protein. The resulting PCR fragment or the insert of BA.3.2 S were subcloned into the KpnI-NotI site of the pCAGGS vector[11] using In-Fusion HD Cloning Kit (Takara, Cat# Z9650N). Nucleotide sequences were determined by DNA sequencing services (Eurofins), and the sequence data were analyzed by SnapGene software v6.1.1 (www.snapgene.com). Pseudoviruses were prepared as previously described.[6-10] Briefly, lentivirus (HIV-1)-based, luciferase-expressing reporter viruses were pseudotyped with the SARS-CoV-2 S. One prior day of transfection, the LentiX-293T cells were seeded at a density of 2 × $10^6$ cells. The LentiX-293T cells were cotransfected with 1 µg psPAX2-IN/HiBiT (a packaging plasmid encoding the HiBiT-tag-fused integrase[1], 1 µg pWPI-Luc2 (a reporter plasmid encoding a firefly luciferase gene[12] and 500 ng plasmids expressing parental S or its derivatives using TransIT-293 transfection reagent (Mirus, Cat# MIR2704) according to the manufacturer's protocol. Two days post transfection, the culture supernatants were harvested and filtrated. The pseudoviruses were harvested and stored at –80°C until use after filtration.

**Neutralization assay**

Neutralization assays were performed previously described[6-10] and mainly conducted by a semi-automated high-throughput method using Fluent780 (Tecan).[7-10,13] The SARS-CoV-2 spike pseudoviruses (counting ~100,000 relative light units) and serially diluted (40-fold to 29,160-fold dilution at the final concentration) heat-inactivated sera were manually prepared in a 2-ml 96-well plate (Greiner, Cat# 780271) and in 96-well microplates (ThermoFisher Scientific, Cat# 168136), respectively. The pseudoviruses were dispensed and mixed with the sera in 384-well plates (ThermoFisher Scientific, Cat# 164610) on Fluent780 (Tecan). Pseudoviruses without sera were included as controls. After incubation at 37°C for 1 hour, HOS-ACE2/TMPRSS2 cells (6,000 cells/30 µl) were added to the 20 µl mixture of pseudovirus and serum in the 384-well white plate on the device. Two days post infection, the infected cells were lysed with a Bright-Glo luciferase assay system (Promega, Cat# E2620) on Fluent780 (Tecan), and the luminescent signal was measured and processed using an Infinite200 and a Magellan (Tecan). The assay of each serum sample was performed in quadruplicate, and the 50% neutralization titer ($NT_{50}$) was calculated using Prism 9 (GraphPad Software).

**Table S1. Human vaccination sera used in this study**

| LP.8.1 vaccine manufacturers | Donor ID | Sex | Age | Date of 1st vaccination (YYYY-MM-DD) | Date of 2nd vaccination (YYYY-MM-DD) | Date of 3rd vaccination (YYYY-MM-DD) | Date of 4th vaccination (YYYY-MM-DD) | Date of 5th vaccination (YYYY-MM-DD) | Date of 6th vaccination (YYYY-MM-DD) | Date of 7th vaccination (YYYY-MM-DD) | Date of 8th vaccination (YYYY-MM-DD) | Date of 9th vaccination (YYYY-MM-DD) | Date of sampling (before vaccination) (YYYY-MM-DD) | Date of LP8.1 vaccination (YYYY-MM-DD) | Date of sampling (after vaccination) (YYYY-MM-DD) | Time interval between vaccination and the second sampling | Prior infection? (YYYY-MM-DD) |
|---|---|---|---|---|---|---|---|---|---|---|---|---|---|---|---|---|---|
| Pfizer/BioNTech | UT0001 | Male | 36 | 2021-07-19M | 2021-08-20M | 2022-05-13M | 2022-11-22MBA.4/5 | 2023-10-01MXBB | 2024-10-03PJN.1 | - | - | - | 2025-10-01 | 2025-10-02 | 2025-10-23 | 21 | Yes (2023-07-03) |
| Pfizer/BioNTech | UT0002 | Female | 56 | 2021-06-26P | 2021-07-17P | 2022-02-05P | 2022-11-16PBA.4/5 | 2024-10-04PJN.1 | - | - | - | - | 2025-09-25 | 2025-10-03 | 2025-10-30 | 27 | No |
| Pfizer/BioNTech | UT0003 | Male | 30 | NA | 2023-2-15MBA.4/5 | 2024-10-03PJN.1 | - | - | - | - | - | - | 2025-09-24 | 2025-09-30 | 2025-10-23 | 23 | No |
| Pfizer/BioNTech | UT0004 | Male | 28 | 2021-07-27M | NA | 2022-07-10P | 2023-01-18PBA.4/5 | 2024-10-03PJN.1 | - | - | - | - | 2025-09-25 | 2025-09-30 | 2025-10-23 | 23 | No |
| Pfizer/BioNTech | UT0005 | Male | 43 | 2021-06-17P | 2021-07-07P | 2022-03-28M | 2022-10-27MBA.4/5 | 2023-09-20PXBB | 2024-10-03PJN.1 | - | - | - | 2025-10-02 | 2025-10-02 | 2025-10-24 | 22 | Yes (2023-06-29) |
| Pfizer/BioNTech | UT0006 | Female | 34 | 2021-03-03P | 2021-8-5P | 2022-2P | 2022-11P | 2023-11P | 2024-10-04PJN.1 | - | - | - | 2025-09-24 | 2025-09-30 | 2025-10-22 | 22 | Yes (2021-06) |
| Pfizer/BioNTech | UT0007 | Male | 32 | 2021-07-28M | 2021-08-25M | 2022-03-10M | 2024-10-03PJN.1 | - | - | - | - | - | 2025-09-25 | 2025-09-29 | 2025-10-23 | 24 | Yes (2022-08-07) |
| Pfizer/BioNTech | UT0008 | Male | 28 | 2021-08-06M | 2021-09-03M | 2024-10-03PJN.1 | - | - | - | - | - | - | 2025-09-24 | 2025-10-02 | 2025-10-23 | 21 | Yes (2022-11-05) |
| Pfizer/BioNTech | UT0009 | Female | 55 | 2021-08-18P | 2021-09-08P | 2022-04-13P | 2022-10-21P | 2024-10-04PJN.1 | - | - | - | - | 2025-09-25 | 2025-10-03 | 2025-10-30 | 27 | Yes (2023-07-24) |
| Pfizer/BioNTech | UT0010 | Male | 41 | 2021-06-28 | 2021-07-19 | 2022-05-17 | 2022-09-03 | 2024-10-03PJN.1 | - | - | - | - | 2025-09-24 | 2025-09-29 | 2025-10-23 | 24 | No |
| Pfizer/BioNTech | UT0011 | Male | 30 | 2021-05-28S | 2021-06-30S | 2021-12-21J | 2024-10-03PJN.1 | - | - | - | - | - | 2025-09-25 | 2025-10-02 | 2025-10-30 | 28 | Yes (2023-05) |
| Pfizer/BioNTech | UT0013 | Female | 29 | 2021-09-12M | 2021-10-10M | 2022-07-19M | - | - | - | - | - | - | 2025-09-24 | 2025-09-29 | 2025-10-24 | 25 | Yes (2023-05-27, 2023-09-11, 2024-08-31) |
| Pfizer/BioNTech | UT0014 | Female | 38 | 2021-07-03M | 2021-07-31M | 2022-02-19M | 2022-12-09P | - | - | - | - | - | 2025-09-24 | 2025-09-29 | 2025-10-23 | 24 | No |
| Pfizer/BioNTech | UT0015 | Male | 31 | 2021-03-07 | 2021-03-28 | 2021-11-20 | NA | - | - | - | - | - | 2025-09-24 | 2025-09-30 | 2025-10-23 | 23 | Yes (2022-12-19) |
| Pfizer/BioNTech | UT0016 | Male | 28 | 2021-8-11P | 2021-09-01P | 2022-04-01M | 2023-10-26PXBB | - | - | - | - | - | 2025-09-24 | 2025-09-30 | 2025-10-22 | 22 | Yes (2022-11-04) |
| Pfizer/BioNTech | UT0017 | Female | 24 | 2021-07-28M | 2021-08-25M | - | - | - | - | - | - | - | 2025-09-25 | 2025-10-03 | 2025-10-30 | 27 | Yes(2021-05-10) |
| Pfizer/BioNTech | UT0018 | Female | 40 | 2021-07-19M | 2021-08-16M | 2022-04-29M | 2022-11-30PBA.4/5 | 2023-12-02PXBB | - | - | - | - | 2025-09-24 | 2025-10-03 | 2025-10-29 | 26 | Yes (2025-03-03) |
| Pfizer/BioNTech | UT0019 | Female | 51 | 2021-07-20P | 2021-08-12P | 2022-02-22M | 2022-11-08PBA.4/5 | - | - | - | - | - | 2025-09-25 | 2025-09-29 | 2025-10-23 | 24 | Yes (2022-11-21) |
| Pfizer/BioNTech | UT0020 | Female | 19 | 2021-09-19P | 2021-10-10P | - | - | - | - | - | - | - | 2025-09-25 | 2025-09-30 | 2025-10-23 | 23 | No |
| Pfizer/BioNTech | UT0021 | Female | 52 | 2021-09-25P | 2021-10-16P | 2022-04-16P | 2022-11-13PBA.4/5 | - | - | - | - | - | 2025-09-25 | 2025-10-02 | 2025-10-30 | 28 | Yes (2024-02-23,2025-07-06) |
| Pfizer/BioNTech | UT0022 | Male | 27 | 2021-06-26J | 2021-12-20P | - | - | - | - | - | - | - | 2025-09-25 | 2025-10-02 | 2025-10-23 | 21 | Yes (2022-04-01) |
| Pfizer/BioNTech | UT0023 | Female | 19 | 2021-09-26 | 2021-10-17 | 2022-06-05 | - | - | - | - | - | - | 2025-09-24 | 2025-10-03 | 2025-10-30 | 27 | Yes(2022-11-29) |
| Pfizer/BioNTech | UT0024 | Female | 25 | 2020-08-12 | 2021-08-20 | 2022-02-26 | - | - | - | - | - | - | 2025-09-25 | 2025-10-03 | 2025-10-30 | 27 | Yes (2023-10-01) |
| Pfizer/BioNTech | 730016 | Female | 36 | NA | 2021 | 2024-10-10PJN.1 | - | - | - | - | - | - | 2025-10-02 | 2025-10-02 | 2025-10-24 | 22 | Yes (2022-12) |
| Pfizer/BioNTech | 730017 | Female | 28 | NA | NA | NA | NA | NA | NA | 2023-10 | 2024-10-10PJN.1 | - | 2025-10-02 | 2025-10-02 | 2025-10-24 | 22 | Yes (2022-09) |
| Pfizer/BioNTech | 730020 | Female | 25 | NA | NA | NA | NA | NA | NA | 2023-08 | 2024-10-10PJN.1 | - | 2025-10-02 | 2025-10-03 | 2025-10-24 | 21 | Yes (2023-08) |
| Pfizer/BioNTech | 730019 | Female | 28 | NA | NA | NA | NA | NA | NA | 2023-10 | 2024-10-10PJN.1 | - | 2025-10-02 | 2025-10-02 | 2025-10-24 | 22 | No |
| Pfizer/BioNTech | 730043 | Female | 29 | NA | NA | 2022-10 | - | - | - | - | - | - | 2025-10-02 | 2025-10-03 | 2025-10-24 | 21 | Yes (2025-02) |
| Pfizer/BioNTech | 730051 | Female | 37 | NA | NA | 2022-10 | - | - | - | - | - | - | 2025-10-02 | 2025-10-02 | 2025-10-24 | 22 | No |
| Novavax/Takeda | S0026 | Male | 30 | 2021-06-24M | 2021-07-22M | 2022-02-01M | 2022-10-04P | 2023-09-20P | 2024-10-01MJN.1 | - | - | - | 2025-09-22 | 2025-09-22 | 2025-10-14 | 22 | Yes (2022-12-01) |
| Novavax/Takeda | S0027 | Male | 57 | 2021-07-30M | 2021-08-27M | 2022-02-27M | 2022-09-22P | 2023-09-20P | 2024-05-17P | 2024-11-18NJN.1 | - | - | 2025-09-22 | 2025-09-22 | 2025-10-14 | 22 | No |
| Novavax/Takeda | S0028 | Male | 41 | 2021-06-22P | 2021-07-13P | 2022-02-23M | 2022-07-29M | 2022-11-30M | 2023-10-04P | 2024-11-02MSPJN.1 | - | - | 2025-09-22 | 2025-09-22 | 2025-10-14 | 22 | No |
| Novavax/Takeda | S0052 | Male | 38 | 2021-08-04M | 2021-09-01M | 2022-03-05M | 2022-09-23P | 2023-10-07M | 2024-11-22MSPJN.1 | - | - | - | 2025-09-22 | 2025-09-22 | 2025-10-14 | 22 | No |
| Novavax/Takeda | S0057 | Female | 41 | 2021-06-30P | 2021-07-29P | 2022-03-03P | 2022-08-19P | 2022-12-02P | 2023-07-14P | 2023-10-31P | 2024-05-29M | 2024-11-06NJN.1 | 2025-09-24 | 2025-09-24 | 2025-10-17 | 23 | No |
| Novavax/Takeda | S0062 | Female | 46 | 2021-08-13P | 2021-09-17P | 2022-03-18M | 2022-10-17P | 2023-10-04M | 2024-10-26MSPJN.1 | - | - | - | 2025-09-19 | 2025-09-19 | 2025-10-14 | 25 | No |
| Novavax/Takeda | S0063 | Female | 50 | 2021-06-21M | 2021-07-21M | 2022-01-31M | 2022-10-01M | 2023-10-04M | 2024-11-09MSPJN.1 | - | - | - | 2025-09-22 | 2025-09-22 | 2025-10-14 | 22 | No |
| Novavax/Takeda | S0065 | Male | 52 | 2021-08-03M | 2021-08-31M | 2022-04-16M | 2022-12-08M | 3023-10-04M | 2024-11-09MSPJN.1 | - | - | - | 2025-09-22 | 2025-09-22 | 2025-10-14 | 22 | No |
| Novavax/Takeda | S0079 | Female | 41 | 2021-07-16M | 2021-08-13M | 2022-03-07M | 2022-10-11P | 2023-07-08P | 2023-11-24M | 2024-06-14M | 2024-11-16MSPJN.1 | - | 2025-09-22 | 2025-09-22 | 2025-10-15 | 23 | Yes (2022-12-16) |
| Novavax/Takeda | S0082 | Female | 47 | 2021-07-19P | 2021-08-09P | 2022-02-12M | 2022-10-14M | 2023-10-07M | 2025-06-26M | 2025-11-02MSPJN.1 | - | - | 2025-09-22 | 2025-09-22 | 2025-10-18 | 26 | No |
| Novavax/Takeda | S0101 | Female | 54 | 2021-06-29P | 2021-07-20P | 2022-02-12M | 2022-10-28M | 2023-09-20P | 2024-11-03MSPJN.1 | - | - | - | 2025-09-19 | 2025-09-19 | 2025-10-14 | 25 | Yes (2024-07-02) |
| Novavax/Takeda | S0102 | Male | 47 | 2021-07-15M | 2021-08-17M | 2022-03-15M | 2022-10-17M | 2023-10-11M | 2024-09-27MJN.1 | - | - | - | 2025-09-22 | 2025-09-22 | 2025-10-17 | 25 | Yes (2025-03-25) |
| Novavax/Takeda | S0153 | Female | 43 | 2021-05-01P | 2021-05-22P | 2022-03-31P | 2022-09-29P | 2023-12-15P | - | - | - | - | 2025-09-19 | 2025-09-19 | 2025-10-15 | 26 | Yes (2022-12-24) |
| Novavax/Takeda | S0165 | Male | 45 | 2021-07-24P | 2021-08-04P | 2022-02-09M | 2022-10-11M | 2023-09-21P | 2024-06-07P | 2024-12-07MSPJN.1 | - | - | 2025-09-22 | 2025-09-22 | 2025-10-20 | 28 | Yes (2025-07-25) |
| Novavax/Takeda | S0173 | Male | 42 | 2021-08-30M | 2021-09-27M | 2022-03-31M | 2022-11-29M | 2023-09-25P | 2024-11-19MSPJN.1 | - | - | - | 2025-09-24 | 2025-09-24 | 2025-10-17 | 23 | No |
| Novavax/Takeda | S0200 | Female | 23 | 2021-07-08M | 2021-08-17M | 2022-04-13M | 2022-11-26P | 2023-09-20P | - | - | - | - | 2025-09-24 | 2025-09-24 | 2025-10-17 | 23 | Yes (2022-03-16) |
| Novavax/Takeda | S0201 | Female | 42 | 2021-07-01M | 2021-07-30M | 2022-02-23M | 2022-10-05M | 2023-12-23D | 2024-06-07P | 2024-11-04MSPJN.1 | - | - | 2025-09-19 | 2025-09-19 | 2025-10-15 | 26 | Yes (2023-09-17) |
| Novavax/Takeda | S0209 | Female | 58 | 2021-07-30P | 2021-08-27P | 2022-03-04P | 2022-10-14M | 2023-10-18N | 2024-11-04MSPJN.1 | - | - | - | 2025-09-24 | 2025-09-24 | 2025-10-15 | 21 | No |
| Novavax/Takeda | S0231 | Female | 46 | 2021-06-25P | 2021-07-30P | 2022-02-21M | 2022-11-25P | 2023-09-29P | 2024-11-15MSPJN.1 | - | - | - | 2025-09-19 | 2025-09-19 | 2025-10-14 | 25 | Yes (2021-01-20) |
| Novavax/Takeda | S0244 | Male | 43 | 2021-07-21M | 2021-08-23M | 2022-03-11M | 2023-12-27P | 2024-07-10M | 2024-11-15MSPJN.1 | - | - | - | 2025-09-24 | 2025-09-24 | 2025-10-14 | 20 | Yes (2022-07-16) |

NA, not applicable.

A, Astrazeneca; P, Pfizer/BioNTech; M, Moderna; S, Sinovac; J, Jenner; N, Novavax/Takeda; MSP, Meji Seika Pharma

BA1/2, BA.1/2 bivalent vaccine; BA4/5, BA.4/5 bivalent vaccine; XBB, XBB.1.5 monovalent vaccine; JN.1, JN.1 monovalent vaccine

**Table S2. Primers used in this study**

| Primer name | Primer sequence (5'-to-3') | Purpose |
|---|---|---|
| Omicron universal Fw | cactatagggcgaattgggtaccatgtttgtgttcctggt | Preparation of S expression plasmid |
| BA.2 WT Rv | agctccaccgcggtggcggccgctcaggtgtagtgcagtttca | Preparation of S expression plasmid |
| S:S31P Fw | caaagctacaccaacCCCttcaccaggggagtc | Preparation of S expression plasmid |
| S:S31P Rv | gactcccctggtgaaGGGgttggtgtagctttg | Preparation of S expression plasmid |
| S:S59F Fw | ctgttcctgccattcTTCagcaatgtgacctgg | Preparation of S expression plasmid |
| S:S59F Rv | ccaggtcacattgctGAAgaatggcaggaacag | Preparation of S expression plasmid |
| S:K182R/S184G/R190S Fw | gacttggagggcAGAcagGGCaacttcaagaacctgAGCgagtttgtgttc | Preparation of S expression plasmid |
| S:K182R/S184G/R190S Rv | gaacacaaactcGCTcaggttcttgaagttGCCctgTCTgccctccaagtc | Preparation of S expression plasmid |
| S:R346T Fw | ggtgttcaatgccaccACCtttgcctctgtctatg | Preparation of S expression plasmid |
| S:R346T Rv | catagacagaggcaaaGGTggtggcattgaacacc | Preparation of S expression plasmid |
| S:S435A/K444R/V445R Fw | ggctgtgtgattGCCtggaacagcaacaagctggacagcAGAAGAagcggcaactac | Preparation of S expression plasmid |
| S:S435A/K444R/V445R Rv | gtagttgccgctTCTTCTgctgtccagcttgttgctgttccaGGCaatcacacagcc | Preparation of S expression plasmid |
| S:T478K/N487D Fw | ccaggctggcaacAAGccatgtaagggaaagggccccGACtgttactttccac | Preparation of S expression plasmid |
| S:T478K/N487D Rv | gtggaaagtaacaGTCggggccctttcccttacatggCTTgttgccagcctgg | Preparation of S expression plasmid |
| S:T572I Fw | gggacattgtggacATCacagatgctgtgag | Preparation of S expression plasmid |
| S:T572I Rv | ctcacagcatctgtGATgtccacaatgtccc | Preparation of S expression plasmid |

**Table S3. Statistics for neutralization titers in 2024 and 2025**

| | Fold change* | p value** | Fold change | p value |
|---|---|---|---|---|
| | Pre vs Post in 2024 | | Pre vs Post in 2025 | |
| B.1.1 | 2.2 | 0.0005 | 1.3 | <0.0001 |
| BA.5 | 4.1 | 0.0005 | 1.6 | <0.0001 |
| XBB.1.5 | 4.7 | 0.0005 | 2.1 | <0.0001 |
| JN.1 | 8.5 | 0.0005 | 2.1 | <0.0001 |
| LP.8.1 | 6.7 | 0.0005 | 3.1 | <0.0001 |
| XEC | 5.8 | 0.001 | 2.8 | <0.0001 |
| NB.1.8.1 | 7.8 | 0.0005 | 2.6 | <0.0001 |
| XFG | 2.6 | 0.0005 | 2.7 | <0.0001 |
| BA.3 | 3.4 | 0.0005 | 1.4 | <0.0001 |
| | Post in 2024 vs Pre in 2025 | | | |
| B.1.1 | 0.4 | 0.0005 | | |
| BA.5 | 0.4 | 0.0009 | | |
| XBB.1.5 | 0.4 | 0.0021 | | |
| JN.1 | 0.3 | 0.0064 | | |
| LP.8.1 | 0.4 | 0.0066 | | |
| XEC | 0.4 | 0.014 | | |
| NB.1.8.1 | 0.3 | 0.0066 | | |
| XFG | 0.6 | 0.013 | | |
| BA.3 | 0.4 | 0.0079 | | |
| | Pre vs Post in 2024 | | Pre vs Post in 2025 | |
| B.1.1 | 0.6 | 0.023 | 0.9 | 0.47 |
| BA.5 | 0.8 | 0.17 | 2.0 | 0.015 |
| XBB.1.5 | 0.7 | 0.0021 | 2.1 | 0.0024 |
| JN.1 | 0.8 | 0.46 | 4.4 | 0.0018 |
| LP.8.1 | 1.0 | 0.33 | 4.0 | 0.0017 |
| XEC | 1.3 | 0.31 | 2.6 | 0.018 |
| NB.1.8.1 | 1.0 | 0.77 | 2.7 | 0.0015 |
| XFG | 1.5 | 0.16 | 1.8 | 0.0051 |
| BA.3 | 0.7 | 0.086 | 1.5 | 0.0054 |

*Fold change is calculated as an average of median of 50% neutralization titers

**P value is determined by two-sided Wilcoxon signed-rank tests

### Consortia

#### The Genotype to Phenotype Japan (G2P-Japan) Consortium

**The Institute of Medical Science, The University of Tokyo, Japan**
Naoko Misawa, Arnon Plianchaisuk, Ziyi Guo, Kaoru Usui, Wilaiporn Saikruang, Spyridon Lytras, Luca Nishimura, Yusuke Kosugi, Shigeru Fujita, Luo Chen, Jarel Elgin M. Tolentino, Wenye Li, Yukun Zhu, Mika Chiba, Shiho Tanaka, Eiko Ogawa, Kaho Okumura, Tsuki Fukuda, Tamaki Yoshihara, Keiko Koizumi, Hiroaki Unno, Mizuho Ota

**Hokkaido University, Japan**
Takasuke Fukuhara, Tomokazu Tamura, Rigel Suzuki, Saori Suzuki, Shuhei Tsujino, Hayato Ito, Hirofumi Sawa, Naganori Nao, Keita Matsuno, Keita Mizuma, Jingshu Li, Izumi Kida, Yume Mimura, Yuma Ohari, Shinya Tanaka, Masumi Tsuda, Lei Wang, Yoshikata Oda, Zannatul Ferdous, Kenji Shishido, Hiromi Mohri, Miki Iida

**Tokyo Metropolitan Institute of Public Health**
Kenji Sadamasu, Kazuhisa Yoshimura, Hiroyuki Asakura, Isao Yoshida, Mami Nagashima

**Tokai University, Japan**
So Nakagawa

**Kyoto University, Japan**
Kotaro Shirakawa, Akifumi Takaori-Kondo, Kazuo Takayama, Rina Hashimoto, Sayaka Deguchi, Yukio Watanabe, Yoshitaka Nakata, Hiroki Futatsusako, Ayaka Sakamoto, Naoko Yasuhara, Takao Hashiguchi, Tateki Suzuki, Kanako Kimura, Jiei Sasaki, Yukari Nakajima, Hisano Yajima

**Hiroshima University, Japan**
Takashi Irie, Ryoko Kawabata

**Kyushu University, Japan**
Kaori Tabata

**Kumamoto University, Japan**
Terumasa Ikeda, Hesham Nasser, Ryo Shimizu, MST Monira Begum, Michael Jonathan, Yuka Mugita, Sharee Leong, Otowa Takahashi, Takamasa Ueno, Chihiro Motozono, Mako Toyoda

**University of Miyazaki, Japan**

Akatsuki Saito, Anon Kosaka, Miki Kawano, Natsumi Matsubara, Tomoko Nishiuchi

**Charles University, Czechia**
Jiri Zahradnik, Prokopios, Andrikopoulos, Miguel Padilla-Blanco, Aditi Konar, Ruojin Tuan

**Acknowledgments**

We would like to thank all members of The Genotype to Phenotype Japan (G2P-Japan) Consortium. We thank Kenzo Tokunaga (National Institute of Infectious Diseases, Japan) for sharing materials, Yuka Kamoshita (Department of Laboratory Medicine, Keio University School of Medicine, Japan), Masayo Noguchi (Clinical Laboratory, Keio University Hospital, Japan) for supporting patient sera collection, and Mika Chiba, Tsuki Fukuda, Mizuho Ota, Tamaki Yoshihara, Keiko Koizumi (Division of Systems Virology, University of Tokyo, Japan) for performing experimental assays.

### Supplementary References


1. Ozono S, Zhang Y, Ode H, et al. SARS-CoV-2 D614G spike mutation increases entry efficiency with enhanced ACE2-binding affinity. *Nat Commun* 2021; **12**(1): 848.
2. Ferreira I, Kemp SA, Datir R, et al. SARS-CoV-2 B.1.617 Mutations L452R and E484Q Are Not Synergistic for Antibody Evasion. *J Infect Dis* 2021; **224**(6): 989-94.
3. Motozono C, Toyoda M, Zahradnik J, et al. SARS-CoV-2 spike L452R variant evades cellular immunity and increases infectivity. *Cell Host Microbe* 2021; **29**(7): 1124-36 e11.
4. Kimura I, Yamasoba D, Tamura T, et al. Virological characteristics of the SARS-CoV-2 Omicron BA.2 subvariants, including BA.4 and BA.5. *Cell* 2022; **185**(21): 3992-4007 e16.
5. Uriu K, Ito J, Zahradnik J, et al. Enhanced transmissibility, infectivity, and immune resistance of the SARS-CoV-2 omicron XBB.1.5 variant. *Lancet Infect Dis* 2023; **23**(3): 280-1.
6. Kosugi Y, Kaku Y, Hinay Jr AA, et al. Antiviral humoral immunity against SARS-CoV-2 omicron subvariants induced by XBB.1.5 monovalent vaccine in infection-naive and XBB-infected individuals. *Lancet Infect Dis* 2024; **24**(3):e147-e148.
7. Kaku Y, Okumura K, Padilla-Blanco M, et al. Virological characteristics of the SARS-CoV-2 JN.1 variant. *Lancet Infect Dis* 2024; **24**(2): e82.
8. Kaku Y, Okumura K, Kawakubo S, et al. Virological characteristics of the SARS-CoV-2 XEC variant. *Lancet Infect Dis* 2024; **24**(12):e736.
9. Chen L, Kaku Y, Okumura K, et al. Virological characteristics of the SARS-CoV-2 LP.8.1 variant. *Lancet Infect Dis* 2025; **25**(4):e193.
10. Uriu K, Okumura K,Uwamino Y, et al. Virological characteristics of the SARS-CoV-2 NB.1.8.1 variant. *Lancet Infect Dis* 2025; **25**(8):e443.
11. Niwa H, Yamamura K, Miyazaki J. Efficient selection for high-expression transfectants with a novel eukaryotic vector. *Gene* 1991; **108**(2): 193-9.
12. Ozono S, Zhang Y, Tobiume M, Kishigami S, Tokunaga K. Super-rapid quantitation of the production of HIV-1 harboring a luminescent peptide tag. *J Biol Chem* 2020; **295**(37): 13023-30.
13. Garcia-Beltran WF, St Denis KJ, Hoelzemer A, et al. mRNA-based COVID-19 vaccine boosters induce neutralizing immunity against SARS-CoV-2 Omicron variant. *Cell* 2022; **185**(3): 457-66 e4.
